# Actin contraction controls nuclear blebbing and rupture independent of actin confinement

**DOI:** 10.1101/2022.12.01.518663

**Authors:** Mai Pho, Yasmin Berrada, Aachal Gunda, Anya Lavallee, Katherine Chiu, Arimita Padam, Marilena L. Currey, Andrew D. Stephens

**Author notes:** Correspondence: Andrew Stephens.

## Abstract

The nucleus is a mechanically stable compartment of the cell that contains the genome and performs many essential functions. Nuclear mechanical components chromatin and lamins maintain nuclear shape, compartmentalization, and function by resisting antagonistic actin contraction and confinement. However, studies have yet to compare chromatin and lamins perturbations side-by-side as well as modulated actin contraction while holding confinement constant. To accomplish this, we used NLS-GFP to measure nuclear shape and rupture in live cells with chromatin decompaction (VPA), loss of lamin B1 (LMNB1-/-), and loss of lamin A/C (LMNA-/-). We then modulated actin contraction while maintaining actin confinement measured by nuclear height. Wild type, chromatin decompaction, and lamin B1 null present bleb-based nuclear deformations and ruptures dependent on actin contraction and independent of actin confinement. Inhibition of actin contraction by Y27632 decreased nuclear blebbing and ruptures to near 0% of cells while activation of actin contraction by CN03 increased the frequency of ruptures by nearly two-fold. However, lamin A/C null results in overall abnormal shape, but similar blebs and ruptures as wild type which were unaffected by actin contraction modulation. Actin contraction control of nuclear shape and ruptures showed that DNA damage levels were more correlated with perturbed nuclear shape than they were with changes in nuclear ruptures. We reveal that lamin B1 is a chromatin decompaction phenotype because using GSK126, which mimics the loss of facultative heterochromatin in lamin B1 null, is sufficient to phenocopy increased nuclear blebbing and ruptures. Furthermore, even though blebs and ruptures in lamin A/C null cells are insensitive to actin contraction, they do have the capacity to form increased levels of nuclear blebs and bleb-based ruptures, shown by treating with VPA. Thus, nuclear bleb formation and bleb-based nuclear ruptures are driven by actin contraction and independent of changes in actin confinement.

## Introduction

The nucleus is the organelle that protects and compartmentalizes the genome and its vital functions that dictate cellular behavior. The nucleus and its main mechanical components chromatin and lamins must properly resist cytoskeletal and/or external forces to maintain the shape and integrity of the nucleus as a compartment (Kalukula *et al*., 2022). Abnormal nuclear morphology is a major hallmark of human disease spanning aging to heart disease to cancer and many others (Butin-Israeli *et al*., 2012; Stephens *et al*., 2019b; Stephens, 2020). In perturbed nuclei, actin running over the top and along the sides of the nucleus antagonizes and causes abnormal nuclear morphology and ruptures (Khatau *et al*., 2009; Le Berre *et al*., 2012; Hatch and Hetzer, 2016). Loss of nucleus compartmentalization via nuclear rupture results in spilling of nuclear contents into the cytoplasm and vice versa (De Vos *et al*., 2011; Vargas *et al*., 2012). Nuclear ruptures have been shown to lead to dysfunction that underlies human disease including causing DNA damage/genomic instability (Denais *et al*., 2016; Irianto *et al*., 2016; Raab *et al*., 2016; Chen *et al*., 2018; Xia *et al*., 2018; Stephens *et al*., 2019a; Shah *et al*., 2021), altering transcription (De Vos *et al*., 2011; Helfand *et al*., 2012), and disruption of cell cycle control (Pfeifer *et al*., 2018). However, it remains unclear what the separate roles of actin contraction vs. actin confinement are in relation to nuclear shape and rupture, respectively the hallmark and driver of human disease.

Actin controls nuclear shape (Khatau *et al*., 2009), but it remains unclear how to separate the effects of actin contraction from actin confinement for both nuclear shape and ruptures. The current dogma is that the actin cytoskeleton compresses the nucleus to cause nuclear blebbing supported by experiments depolymerizing actin which restores shape and compartmentalization. Placing the nucleus under artificial confinement then causes the return of nuclear blebbing and rupture in perturbed nuclei (Le Berre *et al*., 2012; Hatch and Hetzer, 2016). However, in these studies loss of actin disrupts both actin contraction and confinement while artificial confinement induces greater levels of actin confinement than in WT nuclei. Furthermore, pivotal studies of confined migration rely on both confinement as well as actin contraction for migration (Denais *et al*., 2016) because they rely on hormone gradients that both induce directed migration and increase actin contraction (Schneider *et al*., 2009). A recent paper revealed that indeed modulation of actin contraction while under artificial confinement leads to drastic changes in nuclear blebbing and rupture potential (Mistriotis *et al*., 2019). Thus, actin depolymerization and confined migration respectively remove or activate both actin contraction and confinement.

The alternative hypothesis is that actin contraction and not confinement controls nuclear shape and ruptures. Separation of these two roles of actin could be accomplished using a 2D cell culture system (no artificial confinement), modulation of actin contraction, and the ability to measure actin confinement to determine if it changes. Modulation of actin contraction can be accomplished by Rho Activator II, which we denote as CN03 (De Silva *et al*., 2015), and Rho-associated kinase inhibitor Y27632 (Rees *et al*., 2001; Inoue-Mochita *et al*., 2015) which both ultimately alter phosphorylation of myosin light chain 2. To properly assay the independent role of actin contraction in nuclear shape and ruptures, one would need to determine its role across wild type cells and the most reported nuclear perturbations.

The major components of the nucleus responsible for resisting antagonistic actin forces to maintain nuclear shape and compartmentalization include chromatin compaction, lamin B1, and lamin A/C. Perturbations of chromatin structure results in a weaker nucleus, nuclear blebbing, and rupture via altering levels histone modification state (Stephens *et al*., 2018, 2019a; Tamashunas *et al*., 2020; Schibler *et al*., 2022), H1 dynamics (Furusawa *et al*., 2015; Senigagliesi *et al*., 2019), and most recently HP1α (Strom *et al*., 2021) and Hi-C (Belaghzal *et al*., 2021). Lamin B1 loss was one of the first reported perturbations that resulted in nuclear blebbing and rupture (Lammerding *et al*., 2006; Vargas *et al*., 2012). The mechanism remains unclear because different publications have reported conflicting roles of lamin B1 to nuclear mechanics as none (Lammerding *et al*., 2006), stronger in the lamin regime (Shin *et al*., 2013; Stephens *et al*., 2017), and recently weaker (Vahabikashi *et al*., 2022). Finally, perturbation of lamin A/C via knockdown (shRNAi) or knockout (LMNA−/−) is reported to result in reduced nuclear rigidity (Lammerding *et al*., 2006; Pajerowski *et al*., 2007; Swift *et al*., 2013; Stephens *et al*., 2017; Hobson *et al*., 2020) and overall abnormal nuclear shape including lamin A mutants associated with Progeria (Goldman *et al*., 2004; Lammerding *et al*., 2006; Robijns *et al*., 2016; Soria-Valles *et al*., 2016; Stephens *et al*., 2018; Köhler *et al*., 2020). However, depending on perturbation of lamin A/C or if under confined migration the nucleus can also present nuclear blebbing (Goldman *et al*., 2004; Coffinier *et al*., 2010; De Vos *et al*., 2011; Chen *et al*., 2018). Recent work has detailed these different perturbations’ effect on overall cell and nuclear mechanics (Vahabikashi *et al*., 2022). Interestingly, all of these perturbations disrupt chromatin histone modification state, but do not necessarily show the same phenotype. Thus, individual studies focusing on either chromatin decompaction, lamin B1 loss, and lamin A/C loss are known to result in loss of nuclear shape and rupture, but they have yet to be compared side-by-side.

Here we image mouse embryonic fibroblasts with NLS-GFP to track nuclear shape and rupture across multiple nuclear perturbations while modulating actin contraction. Rupture of the nucleus can be assayed by tracking nuclear localization signal green fluorescent protein (NLS-GFP), which provides a diffusible fluorescence marker that accumulates in the nucleus and spills into the cytoplasm upon nuclear envelope rupture. Using NLS-GFP we compare the nuclear shape and rupture behaviors across wild type and nuclear perturbations chromatin decompaction, lamin B1 null cells, and lamin A/C null cells. We establish that actin contraction can be increased (CN03) and decreased (Y27632) without affecting actin confinement in almost all cases. With this approach we find significant changes in nuclear shape and ruptures are driven by actin contraction. However, this change only impacts wild type, chromatin decompaction, and lamin B1 null bleb-based nuclear shape change and ruptures, but not abnormally shaped non-bleb-based lamin A/C null. We reveal that lamin B1 is a chromatin decompaction perturbation while lamin A/C has the capacity to display bleb-based nuclear ruptures, but only when additionally perturbed with VPA. Overall, we provide novel data that actin contraction is essential to nuclear blebbing and rupture while actin confinement is not.

## Results

### A cross comparison of nuclear shape and compartmentalization characteristics in chromatin and lamin perturbations

Loss of chromatin compaction, lamin B1 or lamin A/C are all known to cause abnormal nuclear morphology and rupture, but no studies have done a cross comparison. To accomplish this, we used stable mouse embryonic fibroblasts (MEF) cell lines expressing nuclear localization signal green fluorescent protein (NLS-GFP) to provide live cell tracking over 3 hours at 2-minute intervals. NLS-GFP concentrates in the nucleus to provide nuclear shape measurements and spills into the cytoplasm upon nuclear rupture (**Figure 1A**). Specifically, we scored nuclei as blebbed with > 1 μm diameter protrusion from the main nuclear body and ruptures as a > 25% increase in the NLS-GFP intensity ratio cell/nucleus (**Supplemental Figure 1, movie 1 and 2**; see Materials and Methods). NLS-GFP can then also reaccumulate post nuclear rupture, which reseal on the order of 10 minutes, to provide continual measurements of nuclear shape and the ability to track multiple ruptures for one nucleus (**Supplemental Figure 1 and movie 3**; (Halfmann *et al*., 2019; Young *et al*., 2020)). Using NLS-GFP we tracked nuclear shape, rupture, type of rupture, and frequency of ruptures for wild type (WT) and perturbations of chromatin decompaction (VPA), loss of lamin B1 (LMNB1−/−), and loss of lamin A/C (LMNA−/−).

**Figure 1.**
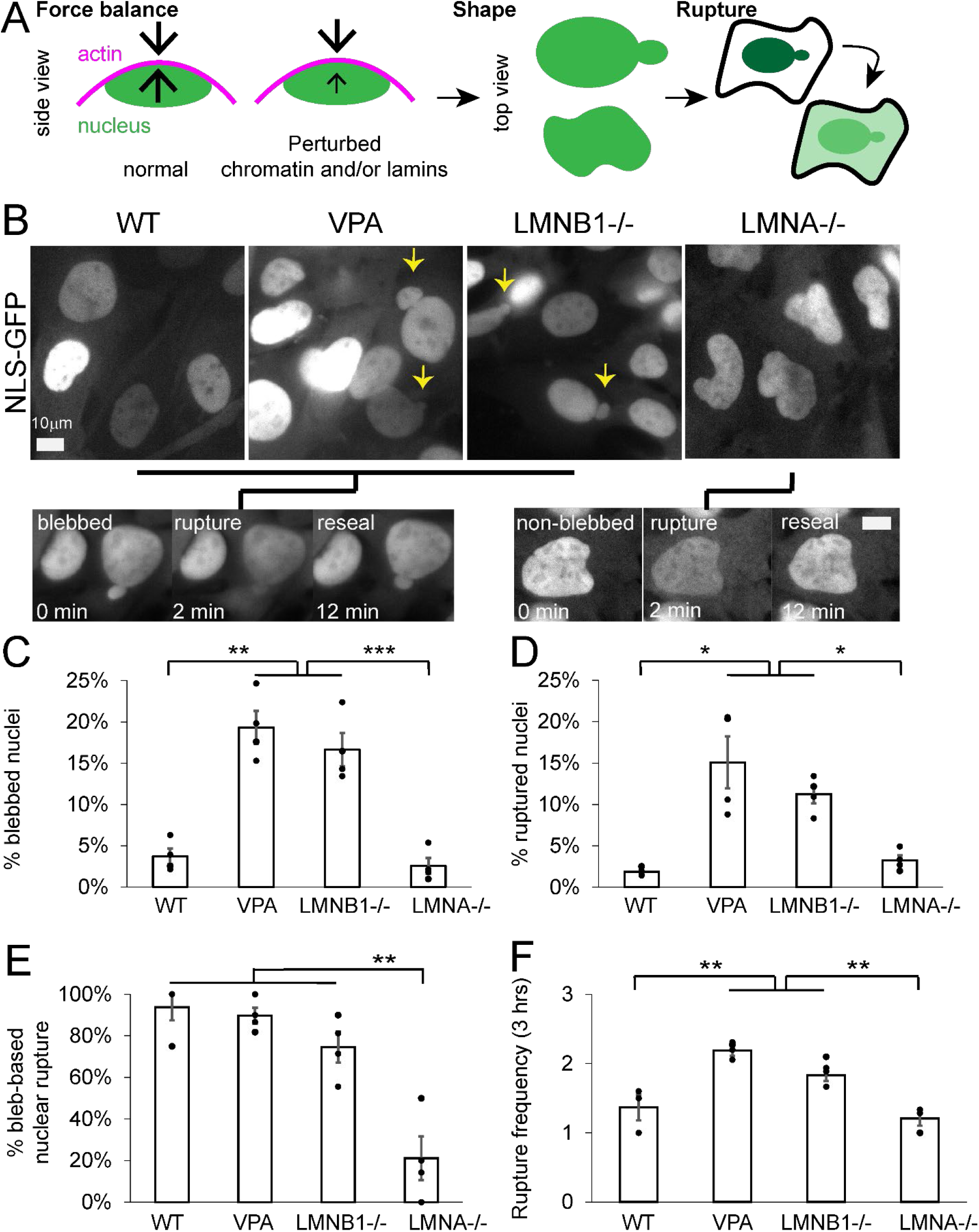
Nuclear morphology and ruptures are bleb-based in WT, VPA, and LMNB1 −/−, but LMNA −/− presents differently. (A) Schematic of force balance between antagonistic actin and resistive nucleus, which can be disrupted by chromatin and/or lamin perturbations leading to abnormal nuclear shape and ruptures. (B) Example images of MEF WT, VPA, LMNB1−/−, and LMNA−/− via NLS-GFP and black lines connecting each to either an example of bleb-based or non-bleb-based nuclear rupture. Graphs of the percentage of nuclei that (C) display a nuclear bleb, (D) rupture at least once, (E) display bleb-based nuclear rupture, and (F) number of nuclear ruptures for a single nucleus that ruptures. All data was measured over a 3-hour period of observation at 2-minute imaging intervals. Each condition has four biological replicates shown as dots. n = WT 102,127,139,158; VPA 227, 146, 170, 108; LMNB1−/− 192, 216, 119, 164; LMNA−/− 111, 142, 149, 101. Student’s t-test p values reported as *<0.05, **<0.01, ***<0.001, no asterisk denotes no significance, p>0.05. Error bars represent standard error. Scale bar = 10 μm.

First, we tracked nuclear blebbing and rupture across all conditions (**Figure 1, B-D**). Wild type nuclei exhibit low levels of nuclear blebbing and rupture at 4 ± 1%. Chromatin decompaction, via histone deacetylase inhibitor VPA, and loss of lamin B1 (LMNB1−/−) both resulted in significantly increased nuclear blebbing (15-20%) and rupture (10-15%) compared to wild type, consistent with previous reports (Lammerding *et al*., 2006; Vargas *et al*., 2012; Hatch and Hetzer, 2016; Chen *et al*., 2018; Stephens *et al*., 2019a). Interestingly, loss of lamin A/C (LMNA−/−) does not result in either an increase in nuclear blebbing or rupture compared to wild type. Instead, LMNA−/− nuclei present as abnormally shaped nuclei that quantitatively measured decreased nuclear circularity (**Supplemental Figure 2**), in agreement with previous reports (Broers *et al*., 2004; Lammerding *et al*., 2006). Thus, loss of nuclear shape via nuclear blebbing correlates with increased nuclear rupture.

To determine if nuclear blebbing could be the main mechanism for nuclear rupture, we tracked nuclear shape upon nuclear rupture. Bleb-based nuclear rupture accounted for > 75% of all nuclear ruptures in wild type, VPA, and LMNB1−/− nuclei (**Figure 1, B and E**). Again, and interestingly, LMNA−/− cells displayed a different behavior where the majority of nuclear ruptures occurred in non-blebbed nuclei at ~ 80% of the time, a complete reversal relative to the other conditions. Thus, nuclear blebbing is the main cause of nuclear rupture for both wild type and most, but not all, nuclear perturbations.

Finally, to determine the frequency of nuclear ruptures from a single nucleus, we tracked number of nuclear ruptures per nucleus that ruptures (**Figure 1F**). The pattern remained consistent with wild type having infrequent multiple nuclear ruptures per nucleus averaging 1.3 ± 0.2. Both VPA and LMNB1−/− nuclei significantly increased the frequency of nuclear ruptures per nucleus to 2.2 ± 0.1 and 1.9 ± 0.1 respectively. LMNA−/− nuclei displayed similar rupture frequency as wild type, 1.2 ± 0.1. Thus, increased nuclear blebbing and bleb-based ruptures upon chromatin decompaction or loss of lamin B1 also results in increased frequency of ruptures in those nuclei.

### Actin contraction modulation independent of actin confinement

Nuclear shape and compartmentalization are determined by the nucleus’ ability to resist antagonistic cytoskeletal and/or external forces working to compress, deform, and rupture the nucleus. Actin contraction is generated by activated myosin motors sliding actin filaments (Murrell *et al*., 2015) and can be measured by phosphorylated myosin light chain 2 (γMLC2, **Figure 2A left**). Actin confinement is due to actin cables running over the top of the nucleus to compress it, which can be measured by nuclear height (**Figure 2A right**). Both actin contraction and confinement can antagonize nuclear shape and stability, but their relative roles remain unknown.

**Figure 2.**
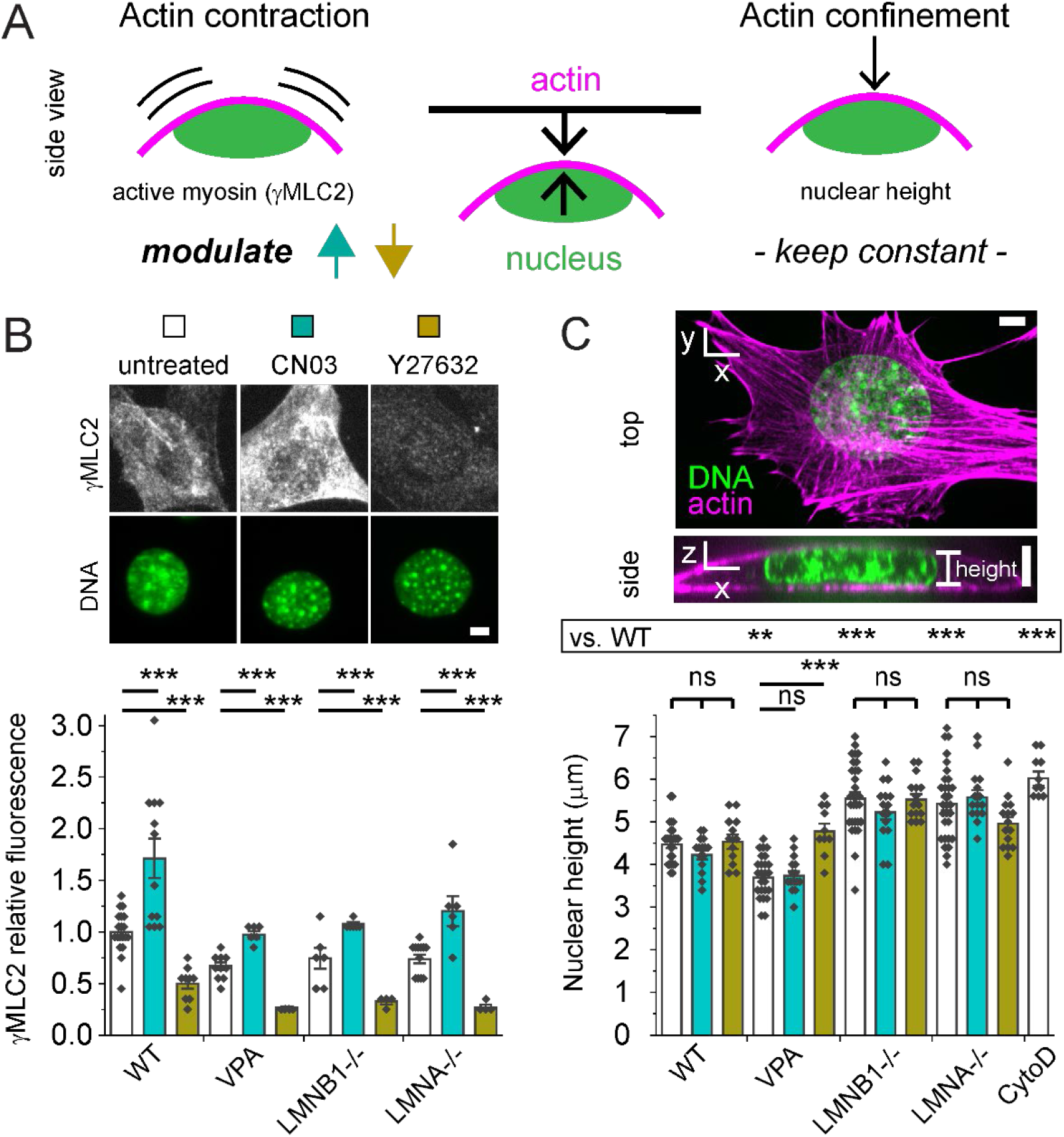
Actin contraction can be modulated while keeping actin confinement constant. (A) Schematic showing that both actin contraction and confinement can antagonize nuclear shape. To separate their effects, we devised an approach to modulate actin contraction while maintaining similar actin confinement. (B) Example images of cells labeled for phosphorylated myosin light chain 2 (γMLC2, white) via immunofluorescence and nuclei stained with Hoechst (DNA, green). Bar graph of relative fluorescence of γMLC2 for WT, VPA, LMNB1−/−, and LMNA−/− without modulation (white bar), with Rho activator II CN03 (turquoise bar), or with ROCK inhibitor Y27632 (gold bar). Biological replicates (diamonds; unmodulated N ≥ 10, CNO3 N ≥ 6, and Y27632 N ≥ 4), each consists of ≥ 20 cells. (C) Representative images of Hoechst/DNA (green) and actin (purple) showing a vertical slice for height and a horizontal maximum projection. Graph of nuclear height measured by Hoechst confocal imaging for WT, VPA, LMNB1−/−, LMNA−/− without modulation (white bar), with Rho activator II CN03 (turquoise bar), or with ROCK inhibitor Y27632 (gold bar). Unmodulated n = 25-30; CNO3 n = 15, Y27632 n = 10-15. WT treated with cytochalasin D (CytoD, actin depolymerization, n = 10) serves as a known control for increase in height. Student’s t-test p values reported as *<0.05, **<0.01, ***<0.001, or ns denotes no significance, p>0.05. Error bars represent standard error. Scale bar = 5 μm.

To determine if actin contraction and/or actin confinement are partially responsible for changes in nuclear shape and rupture upon nuclear perturbations, we tracked both in wild type, VPA, LMNB1−/−, and LMNA−/−. First, we measured actin contraction levels by immunofluorescence of γMLC2. All nuclear perturbations measured a similar level of γMLC2, suggesting changes in nuclear shape via chromatin and lamin perturbations do not arise from changes in actin contraction (**Figure 2B**). Next, we measured nuclear height (**Figure 2C**) which provides a measure of actin confinement and the force balance between the resistive nucleus and the actin that runs over the top of the nucleus compressing it. Nuclear height decreased upon chromatin decompaction via VPA, as expected since the nucleus is softer (Krause *et al*., 2013; Shimamoto *et al*., 2017; Stephens *et al*., 2017; Hobson *et al*., 2020). However, LMNB1−/− and LMNA−/− nuclei, both reported to be softer as well (Vahabikashi *et al*., 2022), did not decrease in nuclear height but instead increased in nuclear height relative to wild type (**Figure 2C**). This data suggests that actin confinement is not a driver of nuclear shape change in loss of either lamin B1 or lamin A/C. Measurements of nuclear height to assess actin contraction in normal vs. blebbed LMNB1−/− nuclei reveal no change, further supporting the idea that actin confinement changes are not essential to nuclear blebbing (**Supplemental Figure 3A**). Overall, nuclear perturbations to chromatin or lamins do not impact actin contraction or confinement in a consistent manner.

While actin contraction stays constant in nuclear perturbations, we hypothesized that it may have an important role in antagonizing nuclear shape and compartmentalization. To modulate actin contraction, we used drugs previously established to affect the Rho pathway via activator CN03 (De Silva *et al*., 2015) and Rho-associated kinase inhibitor Y27632 (Rees *et al*., 2001; Inoue-Mochita *et al*., 2015). Immunofluorescence measurements of γMLC2 confirm across wild type and all nuclear perturbations in our study that activator CN03 significantly increases actin contraction while inhibitor Y27632 drastically decreases it (**Figure 2B**), in agreement with other reports (Hernandez *et al*., 2020). Thus, we established methodology to modulate actin contraction to study its effect on nuclear shape and ruptures.

Alterations in actin contraction could also affect actin confinement. To determine if actin confinement changes upon modulation of action contraction via activator CN03 and inhibitor Y27632, we measured nuclear height in all conditions. In wild type and all nuclear perturbations, except VPA Y27632, nuclear height did not change significantly upon increased or decreased actin contraction (p > 0.05, **Figure 2C**). Thus, actin contraction in most cases can be modulated independently of actin confinement. This data supports an approach that allows us to determine the role of actin contraction in antagonizing nuclear shape and stability.

### Actin contraction is an essential determinant of bleb-based nuclear shape deformations and ruptures

To determine the effect of actin contraction in nuclear blebbing and rupture, we first increased actin contraction via CN03 across all conditions and imaged NLS-GFP for 3 hours at 2-minute intervals. Both wild type and lamin B1 null treated with CN03 to increase actin contraction resulted in both increased percentage of nuclei that blebbed and ruptured (teal, **Figure 3, A and B**). However, chromatin decompaction via VPA showed no change in either blebbing or rupture upon activation of actin contraction via CN03. Tracking nuclear shape in which nuclei ruptured reveals no change in the base behavior of each condition (bleb-based vs. non-bleb-based rupture, **Figure 3C**). Finally, CN03 increased the frequency of ruptures for a single nucleus for all bleb-based nuclear rupture conditions WT, VPA, LMNB1−/− (**Figure 3D)**. This is interesting because while CN03 did not increase the number of nuclei that blebbed or ruptured for VPA, it does nearly double the number of ruptures for a single nucleus that does rupture over a 3 hour period of time. Again, non-bleb-based LMNA−/− showed no change upon CN03 treatment, suggesting it was insensitive. Overall, increased actin contraction via CN03 can increase nuclear blebbing, rupture, and frequency of ruptures for perturbations presenting bleb-based nuclear rupture phenotype.

**Figure 3.**
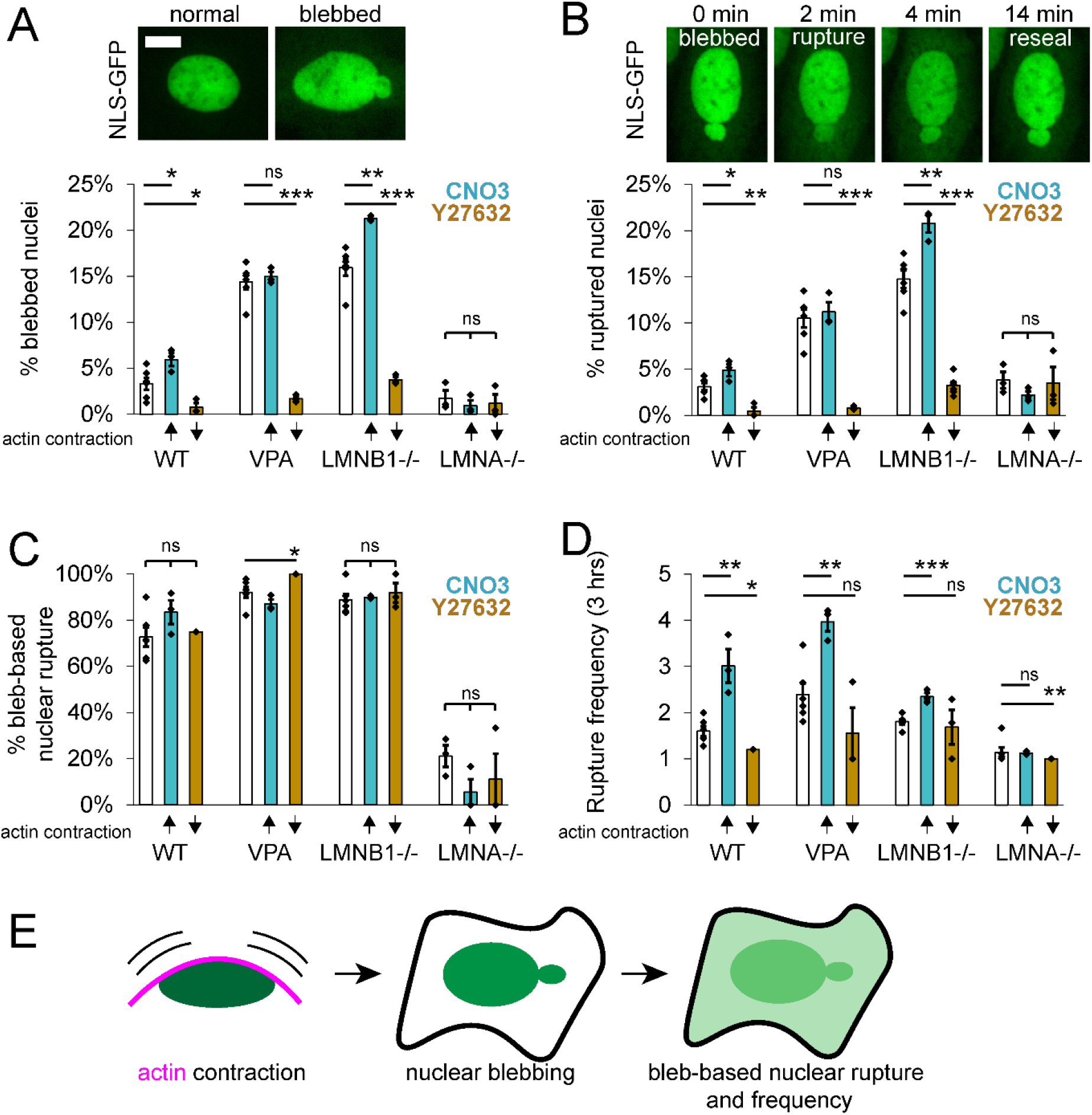
Actin contraction controls nuclear blebbing and bleb-based ruptures, while having little effect on LMNA −/− behaviors. Example images of (A) normal and blebbed nuclei as well as (B) bleb-based nuclear rupture via NLS-GFP time lapse imaging. (A, B, C, D) Graphs showing the percentage of (A) blebbed nuclei, (B) nuclear ruptures, and (C) bleb-based ruptures along with (D) the number of ruptures (frequency) of a single nucleus that ruptures over 3 hours imaged at 2-minute intervals for WT, VPA, LMNB1−/−, LMNA−/− without modulation (white bar), with increased actin contraction (turquoise bar, CN03), or with decreased actin contraction (gold bar, Y27632). 6 biological replicates for unmodulated and 3 biological replicates for increased or decreased actin contraction, experiments represented by black dots, n = 75-400 cells each experiment). Student’s t-test p values reported as *<0.05, **<0.01, ***<0.001, or ns denotes no significance, p>0.05. Error bars represent standard error. Scale bar = 10 μm. (E) Schematic showing that actin contraction leads to nuclear bleb formation and bleb-based nuclear ruptures, where levels of actin contraction determine frequency of nuclear ruptures.

Next, we determined the effects of actin contraction inhibition on nuclear blebbing and ruptures in wild type and nuclear perturbations. Decreased actin contraction via Y27632 revealed a drastic and consistent response decreasing the percentage of nuclei that presented blebs and ruptures for bleb-based conditions WT, VPA, and LMNB1−/− (gold, Y27632, **Figure 3, A and B**). Even wild type’s low levels of nuclear blebbing and ruptures decreased significantly from ~3% to < 0.5% upon actin contraction inhibition, suggesting it is essential for these behaviors. Both VPA and LMNB1−/− perturbations showed a similar nearly essential need for actin contraction to form blebs and cause ruptures as these dropped from 10-20% to < 1-3%. However, LMNA−/− cells’ low percentages of nuclear blebs and ruptures remained unchanged upon decreased actin contraction, continuing the trend of insensitivity to changes in actin contraction. However, nuclear shape measured by circularity in LMNA−/− was significantly improved upon actin contraction inhibition (**Supplemental Figure 2**), showing that LMNA−/− is not completely insensitive. The type of rupture, bleb-based or non, remained similar as well as number of ruptures per nucleus that ruptures (**Figure 3, C and D**), with minor exceptions. Thus, bleb-based nuclear rupture conditions wild type, chromatin decompaction, and lamin B1 null require actin contraction for nuclear blebbing and rupture.

In summary, we find that actin contraction is essential for the bleb-based nuclear rupture phenotype (**Figure 3E**). Furthermore, increased actin contraction can drive more cells to bleb and rupture while driving the number of ruptures for a single nucleus higher as well. Oppositely, actin contraction has little effect on the percentage of nuclei that bleb and rupture in the non-blebbed rupture phenotype of LMNA−/−. Taken together this data supports that, independent of actin confinement, actin contraction is a major determinant of bleb-based nuclear shape and rupture.

### Blebbed or abnormal nuclei present higher levels of DNA damage

Loss of nuclear shape and compartmentalization causes nuclear dysfunction. One of the most common measures of nuclear dysfunction upon abnormal nuclear deformations and ruptures is DNA damage. Thus, we tracked DNA damage via γH2AX foci relative to nuclear shape in WT, VPA, LMNB1−/−, and LMNA−/−. To determine the number of DNA damage foci in a nucleus, we experimentally measured the Gaussian full width half maximum of a diffraction limited focus and empirically determined intensity threshold (see Materials and Methods). Relative to normally shaped nuclei, blebbed nuclei in WT, VPA, and LMNB1−/− displayed greater than a two-fold increase in DNA damage on average (**Figure 4, A and B**), in agreement with previous work (Stephens *et al*., 2019a). LMNA−/− abnormally shaped nuclei, determined by circularity < 0.9, displayed a similar outcome as blebbed nuclei with a significant increase in DNA damage relative to normally shaped nuclei. Thus, across wild type and different nuclear perturbations deformed nuclei present drastically increased levels of DNA damage.

**Figure 4.**
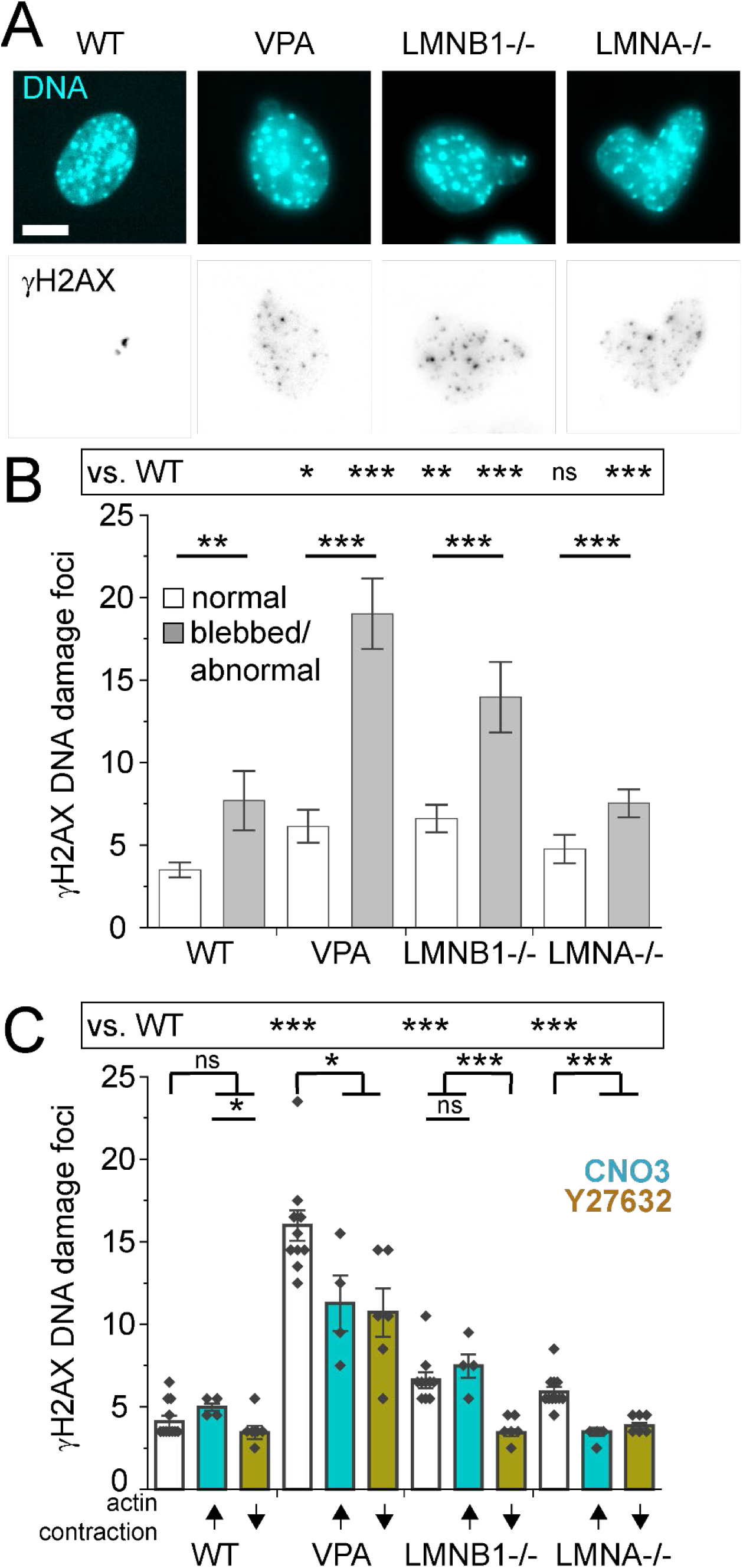
Increased DNA damage is associated with nuclear shape more so than actin contraction. (A) Representative images of nucleus shape via labeling DNA (Hoechst, Cyan) and DNA damage foci (γH2AX, inverted gray scale). (B) Graph of number of γH2AX DNA damage foci for normally shaped nuclei vs. blebbed/abnormal (WT normal n = 115, blebbed n = 17; VPA normal n = 119, blebbed n = 55; LMNB1−/− normal n = 103, blebbed n = 32, LMNA−/− normal n = 73, abnormal circularity < 0.9 n = 214 (see **Supplemental Table 1**). (C) Graph of number of γH2AX DNA damage foci for WT, VPA, LMNB1−/−, LMNA−/− without modulation (white bar), with increased actin contraction (turquoise bar, CN03), or with decreased actin contraction (gold bar, Y27632). Multiple biological replicates of unmodulated (N=10), increased (N=4) and decreased (N=6) actin contraction. Experiments represented by black dots, where n > 40 cells per experiment. Student’s t-test p values reported as *<0.05, **<0.01, ***<0.001, or ns denotes no significance, p>0.05. Error bars represent standard error. Scale bar = 10 μm.

To determine if actin contraction-based changes significantly impact DNA damage levels, we measured the number of γH2AX foci upon activation and inhibition of actin contraction. Our data supports that actin contraction activation via CN03 increases how many times a single nucleus ruptures in bleb-based rupture phenotypes (see **Figure 3D**). Thus, we reasoned that increasing actin contraction would provide a measure of the importance of nuclear ruptures to DNA damage levels. DNA damage measured by γH2AX foci did not significantly increase upon CN03 activation of actin contraction, suggesting the frequency of nuclear ruptures might be less important than abnormal nuclear shape (**Figure 4, C and B,** CN03 vs. abnormal shape, respectively). Treatment of all conditions with actin contraction inhibitor Y27632 significantly decreased DNA damage levels (gold, **Figure 4C**). These results might be expected given that Y27632 decreases nuclear blebbing and abnormal nuclear morphology across all conditions (**Supplemental Figure 2**), recapitulating decreased DNA damage in normally shaped nuclei. This data does not support nuclear ruptures as main mechanism for increased DNA damage but instead suggests abnormal shape as a major driver of DNA damage. Overall, actin contraction driven antagonism of nuclear shape results in nuclear dysfunction measured by increased DNA damage.

### LMNB1−/− loss of heterochromatin is sufficient to generate nuclear blebs and ruptures

We hypothesized that the similarities between VPA and LMNB1−/− might be due to the shared perturbation of changes in histone modification state. While VPA treatment does not cause lamin B1 loss (Stephens *et al*., 2018), it has been reported by many that LMNB1−/− nuclei display chromatin decompaction via loss of facultative heterochromatin (Camps *et al*., 2014; Stephens *et al*., 2018; Vahabikashi *et al*., 2022). To determine if LMNB1−/− nuclei bleb-based phenotype could be due to facultative heterochromatin loss, we recapitulated this facultative heterochromatin loss via treatment with EZH2 methyltransferase inhibitor GSK126 (McCabe *et al*., 2012). Immunofluorescence measurements confirm a significant decrease of facultative heterochromatin marker H3K27me^3^ in LMNB1−/− nuclei relative to wild type (53±3% loss, **Figure 5A**). Treatment with GSK126 revealed a significant decrease in H3K27me^3^ on the order of LMNB1−/− (39±2% loss, **Figure 5A**). Thus, GSK126 treatment roughly recapitulates LMNB1−/− loss of facultative heterochromatin.

**Figure 5.**
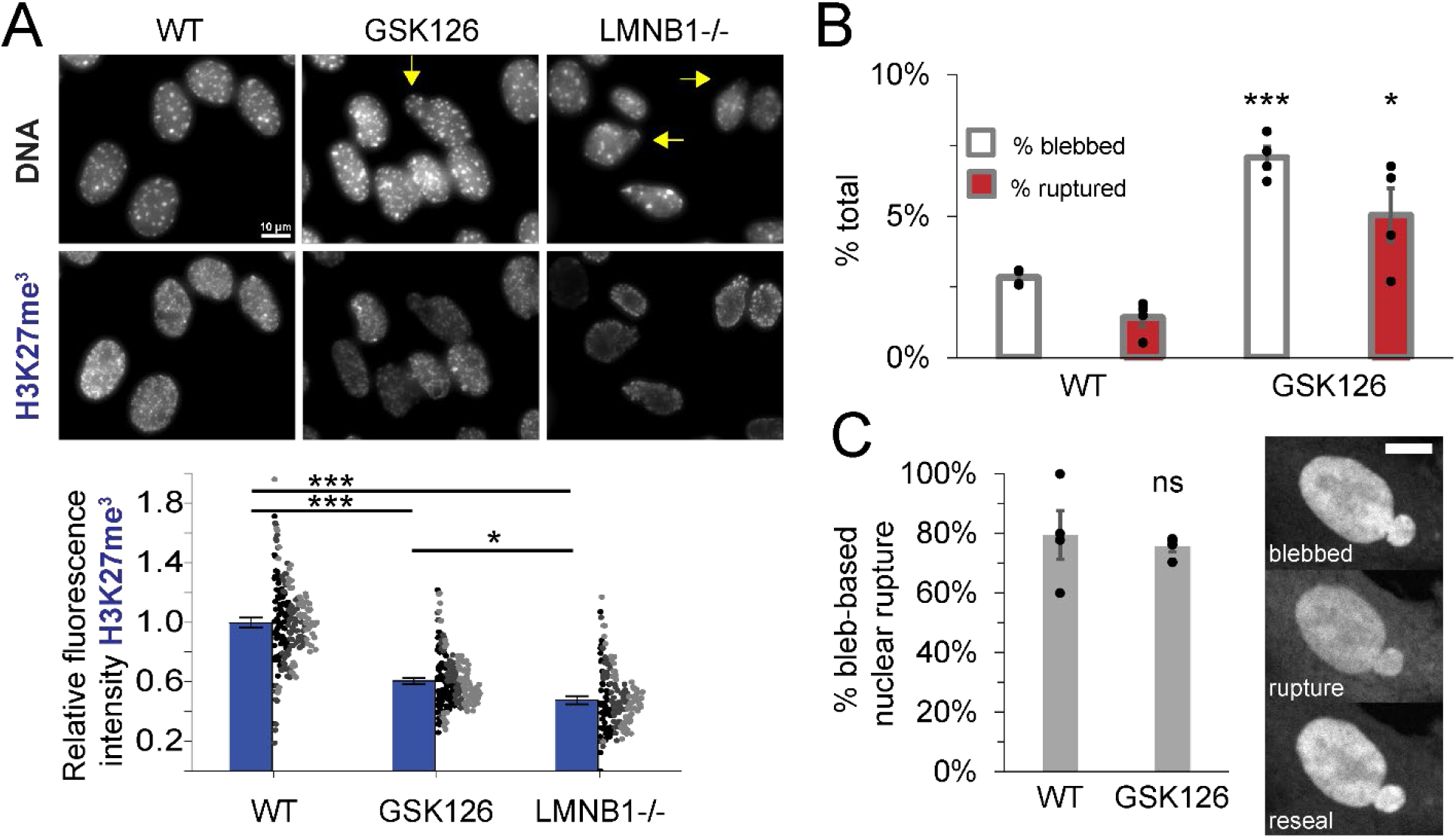
Recapitulating LMNB1−/− loss of facultative heterochromatin alone is sufficient to phenocopy increased nuclear blebbing and rupture. (A) Representative images of nuclei stained for DNA via Hoechst and facultative heterochromatin via H3K27me3 (Blue). Yellow arrow denotes blebbed nucleus. Graph of relative fluorescence intensity of H3K27me3 for wild type (WT), EZH2 inhibitor (GSK126), and lamin B1 null (LMNB1−/−). Dots are individual measures n > 50 nuclei per experiment from three biological replicates, denoted by color (black, dark gray, light gray), were averaged (blue bar). (B) Graph of % total of cells displaying nuclear blebs (white) or ruptures (red) for wild type and GSK 126. (C) Graph of % bleb-based nuclear ruptures for wild type and GSK 126. Four biological replicates displayed as dots with > 250 cells each. Blebbing and ruptures in GSK126 are dependent on actin contraction as Y27632 decreases blebbing to 1% and ruptures to 0% (GSK126 + Y27632 n = 226 cells, p > .01, **Supplemental Table 1**), data not graphed. Student’s t-test p values reported as *<0.05, **<0.01, ***<0.001, no asterisk or ns denotes no significance, p>0.05. Error bars represent standard error. Scale bar = 10 μm.

To determine if loss of facultative heterochromatin is sufficient to induce the bleb-based phenotype of LMNB1−/−, we measured nuclear shape and ruptures in GSK126-treated nuclei. NLS-GFP nuclear shape and rupture tracking in GSK126 treated cells reveals a significant increase in nuclear blebbing and nuclear ruptures compared to wild type (**Figure 5B**). Nuclear ruptures were also found to be bleb-based in the majority of cases similar to wild type (> 80%, **Figure 5C**). Actin contraction was essential for nuclear bleb formation in GSK126-treated cells, as treatment with inhibitor Y27632 decreased nuclear blebbing and rupture to 1% or less (**Supplemental Table 1**). Overall, this data suggests that most of LMNB1−/− nuclear phenotype could be due to loss of facultative heterochromatin. Thus, our data supports that VPA and LMNB1−/− show similar nuclear behaviors because both perturbations are based in histone modification changes that decompact chromatin.

### LMNA−/− are capable of bleb-based deformation and ruptures

We hypothesized that LMNA−/− nuclei do not show bleb-based behaviors because this perturbation cannot, due to reported disrupted nuclear-actin connections (Broers *et al*., 2004; Vahabikashi *et al*., 2022). To determine if LMNA−/− nuclei have the capacity to form nuclear blebs and ruptures, we treated non-bleb-based LMNA−/− with VPA, a condition that causes bleb-based nuclear shape change and rupture. Upon VPA treatment of LMNA−/− MEF cells nuclear blebbing, ruptures, % bleb-based ruptures, and rupture frequency all significantly increased relative to LMNA−/− untreated (**Figure 6**). Interestingly, this increase in bleb-based behaviors was not reliant on changes in actin confinement, as nuclear height did not decrease in LMNA−/− with VPA treatment (**Supplemental Figure 3B**), further suggesting changes in actin confinement are unimportant to nuclear blebbing. Thus, LMNA−/− nuclei have the capacity to display increased nuclear blebbing and ruptures and suggests a more complex reason for why this perturbation behaves differently.

**Figure 6.**
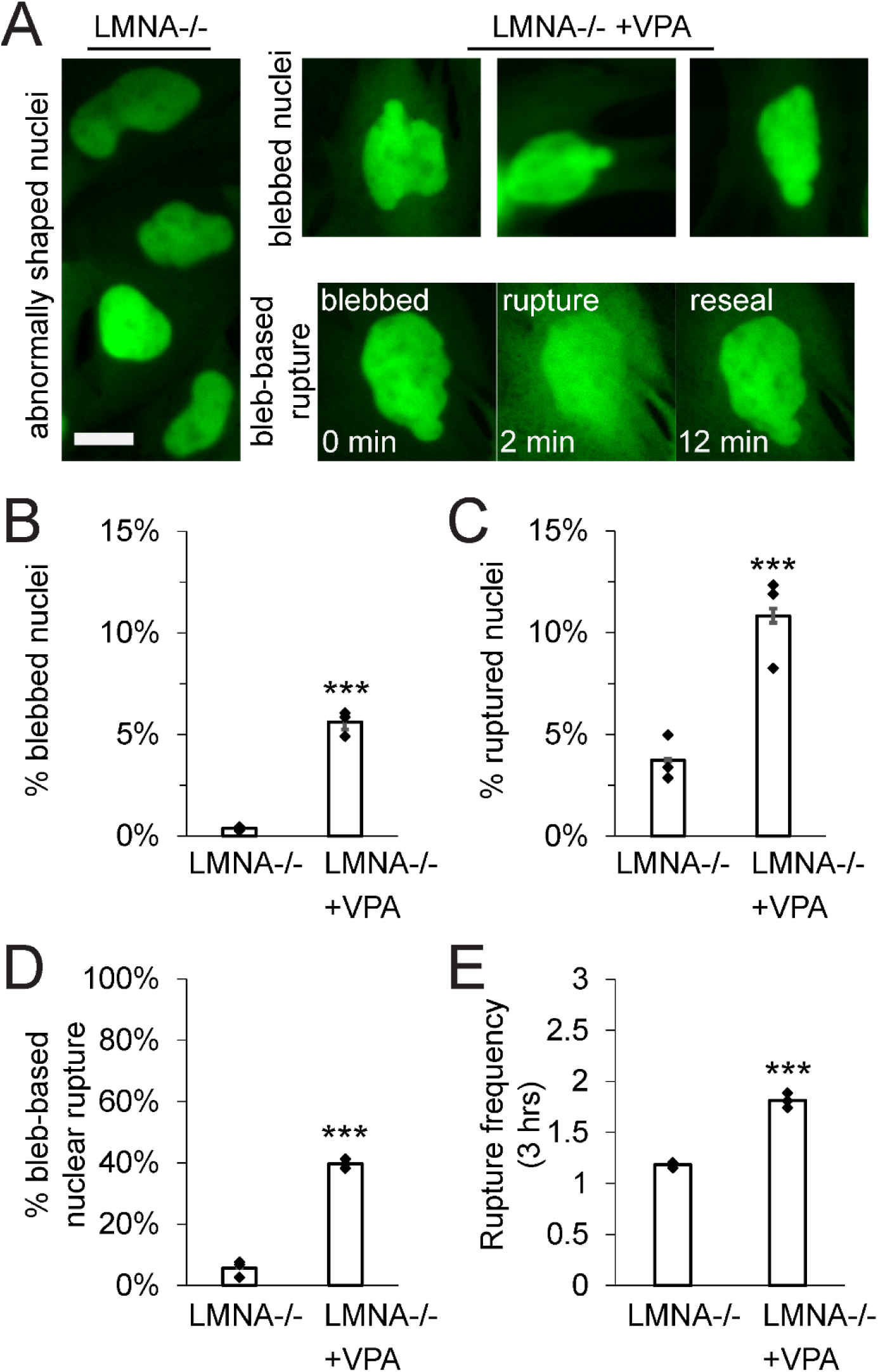
LMNA −/− nuclei have the capacity to form blebs demonstrated by treatment with VPA. (A) Representative images of LMNA−/− abnormally shaped nuclei (left) and LMNA−/− nuclei treated with VPA that display nuclear blebs and bleb-based rupture (right). Graphs of (B) percentage of blebbed nuclei, (C) percentage of ruptured nuclei, (D) percentage of total ruptures that were bleb-based, and (E) nuclear rupture frequency for LMNA−/− and LMNA−/− + VPA imaged for 3 hours at 2-minute intervals via NLS-GFP. Three biological replicates n ≥ 100 cells each for LMNA−/− without or with VPA. Student’s t-test p values report significance *** < 0.001. Error bars represent standard error. Scale bar = 10 μm.

## Discussion

Maintenance of nuclear shape and compartmentalization is determined by a balance between nuclear components resisting deformations induced by the cytoskeleton and external forces. Here we provide novel evidence that actin contraction independent of actin confinement controls nuclear blebbing and ruptures in both wild type and many prominent nuclear perturbations seen in human disease. Through modulation of actin contraction, we show in almost all cases that actin confinement, measured by nuclear height, is unchanged. Inhibition of actin contraction reveals it is essential for nuclear blebbing and ruptures, while activation of actin contraction drastically increased nuclear rupture frequency of single nuclei. Interestingly, actin contraction has less effect on LMNA−/− non-bleb-based behavior but is partially responsible for abnormal nuclear shape in this perturbation. We go on to show that the similarities between VPA and LMNB1−/− are both due to underlying histone modification changes that lead to decompact chromatin. On the other hand, our data show that LMNA−/− nuclei have the capacity to present nuclear blebbing and bleb-based ruptures, revealing that this perturbation will require future studies to further understand its different and complex phenotype. Overall, our work reveals that actin contraction is a major determinant of nuclear blebbing and bleb-based ruptures.

### Actin contraction not confinement controls nuclear shape and ruptures

The previous dogma was that actin fibers running over the top of the nucleus control nuclear shape and compartmentalization via confinement/compression. Pivotal studies of nuclear shape determination reported that actin was a major antagonistic factor to nuclear shape. Loss of nuclear shape overall in progeria cells or nuclear blebs in lamin B1 null cells could be alleviated by removing actin via actin depolymerization drugs such as latrunculin A/B and cytochalasin D (Le Berre *et al*., 2012; Hatch and Hetzer, 2016). In these actin depolymerization treatments, nuclear height was reported to increase since actin was no longer compressing the nucleus, which provided data supporting that actin confinement might control nuclear shape. Abnormal deformations could then be re-established if actin depolymerizers were removed and actin reformed on top of the nucleus causing actin confinement/compression. To prove that actin confinement was responsible, cells with depolymerized actin were compressed via glass plate on top of the cell to restore artificial confinement which resulted in abnormal shape, nuclear blebbing, and nuclear ruptures. These experiments have two weaknesses. First, use of drugs that cause actin depolymerization disrupts both actin contraction as well as confinement. Second, artificial confinement studies over-compress the nucleus significantly more than compared to the nucleus before actin depolymerization. Over-compression of the nucleus causes material failure, which agrees with experiments where nuclei migrate between narrow channels resulting in deformation and rupture (Denais *et al*., 2016; Raab *et al*., 2016), but it does not support that confinement is the major determinant. In order to truly test the relative roles of actin contraction vs. confinement, a different approach emerged.

Our work and others show that actin confinement is not the major mechanism of abnormal nuclear shape and ruptures. Increased nuclear blebbing and ruptures in artificially confined cells might actually be dependent on actin contraction (see discussion below (Mistriotis *et al*., 2019)). Similarly, in unconfined cells our findings show that actin confinement matters very little to nuclear shape, blebbing, and ruptures in nuclear perturbations (**Figure 2**). In contradiction with the current view, actin confinement is less (increased nuclear height) in both LMNB1−/−, which displays increased nuclear blebbing and ruptures, and LMNA−/−, which displays decreased nuclear circularity. Furthermore, in the one perturbation where actin confinement increases (VPA), this change is dependent on actin contraction. Thus, actin contraction might control actin confinement, in agreement with artificial confinement studies (Mistriotis *et al*., 2019). Furthermore, nuclear height remained similar in normally shaped nuclei vs. blebbed nuclei (**Supplemental Figure 3A**). Finally, increased nuclear blebbing and ruptures in LMNA−/− and VPA dual perturbation did not change actin confinement (nuclear height) relative to LMNA−/− (**Supplemental Figure 3B**). Thus, our data and others clearly show how actin confinement is not the main antagonistic factor working to deform and rupture the nucleus in chromatin or lamin perturbations.

Actin contraction is the major antagonistic factor of nuclear shape and compartmentalization. Actin contraction inhibition resulted in a near total loss of nuclear blebs and ruptures in all bleb-based conditions (**Figure 3**). Specifically, already low levels of wild type blebbing and ruptures significantly decreased to a fraction of a percent, strongly suggesting action contraction is the reason any blebbing or ruptures occur in wild type. Our data agrees nicely with work in an artificial confinement system where actin contraction inhibition suppresses nuclear blebbing back to unconfined levels (Mistriotis *et al*., 2019) and that actin contraction causes nuclear strain (Alam *et al*., 2015). Recent work shows that transcription is also a major contributor to nuclear blebbing and ruptures in VPA and LMNB1−/− (Berg *et al*., 2022). However, transcription had no effect on wild type nuclear blebbing percentage, further supporting actin contraction as the main determinant. Finally, increased contraction showed increased nuclear rupture frequency with the capability to increase nuclear blebbing and ruptures in most bleb-based conditions. This again agrees with studies showing increased actin contraction in cells under confinement increases nuclear blebbing (Mistriotis *et al*., 2019). Overall, the data strongly support that actin contraction is the main antagonistic force deforming and rupturing the nucleus.

### Nuclear blebbing is the base phenotype for MEF cells

While many papers have studied the effects of chromatin- or lamin-based perturbations on nuclear shape and ruptures, most have only focused on one perturbation. Excitingly, many current gene screens have provided some measure of nuclear characteristics across many chromatin and lamins protein knockdowns and/or drug treatments (Tamashunas *et al*., 2020; Schibler *et al*., 2022). Other studies have worked to extensively characterize the loss of different lamins (Chen *et al*., 2018; Vahabikashi *et al*., 2022). Here we provide one of the first detailed studies of both a direct chromatin perturbation and lamin perturbations. Interestingly, we find that nuclear bleb formation and bleb-based nuclear ruptures are the dominant form of nuclear shape and compartmentalization loss, at least for MEF cells. These findings are supported by many other studies of individual perturbations but specifically provided for the first time the ability to compare across both chromatin and lamin perturbations.

Chromatin decompaction is now well-known to lead to abnormal nuclear morphologies and ruptures (Stephens *et al*., 2019b). Chromatin decompaction by histone modification alterations or other proteins provides a perturbation of chromatin without altering lamin levels (Furusawa *et al*.,2015; Stephens *et al*., 2018; Strom *et al*., 2021). Oppositely, almost all lamin perturbations, but specifically LMNB1−/− and LMNA−/−, have a secondary effect of altered chromatin, usually through change to histone modifications (Stephens *et al*., 2018, 2019a; Vahabikashi *et al*., 2022). Our data here, provide insight that chromatin decompaction simply exacerbates the wild type phenotype of nuclear blebbing and majority bleb-based ruptures, which both can be almost fully suppressed by an actin contractility inhibitor Y27632 (**Figures 1 and 3**).

Loss of lamin B1 was one of the first and most studied nuclear blebbing perturbations (Lammerding *et al*., 2006; Shimi *et al*., 2008; Vargas *et al*., 2012). More recently, many have shown that loss of lamin B1 results in loss of facultative heterochromatin (Camps *et al*., 2014; Stephens *et al*., 2018, 2019a; Chang *et al*., 2022; Vahabikashi *et al*., 2022), which we recapitulated (**Figure 5A**). Our data reveal that simply decreasing levels of facultative heterochromatin via methyltransferase inhibitor GSK126 is sufficient to increase nuclear blebbing and rupture (**Figure 5, B and C**). This new data agrees with our previous data showing that increased heterochromatin levels via histone demethylase inhibition by methylstat treatment (Stephens *et al*., 2018) and mechanotransduction (Stephens *et al*., 2019a) rescues nuclear shape in LMNB1−/− nuclei. However, loss of facultative heterochromatin alone does not match the high levels of nuclear blebbing and rupture in LMNB1−/−, suggesting loss of lamin B1 causes an additive effect. To ultimately determine if lamin B1 is a chromatin perturbation, we would need to conduct micromanipulation force measurements to assay the separate short-extension chromatin-based regime vs. long-extension lamin-based regime (Stephens *et al*., 2017; Currey *et al*., 2022). Lamin B1’s role in nuclear mechanics, morphology, and compartmentalization remains to be settled, but our data clearly provide evidence that changes to histone modification state are a major contributor to its phenotype.

Lamin A/C loss was the outlier in this group of nuclear perturbations because it presents with a non-bleb-based loss of nuclear shape and compartmentalization. This finding is somewhat confusing as it is well-reported that loss of lamin A/C already causes loss of heterochromatin, a condition shown to cause bleb-based behavior. Our finding of loss of overall shape in LMNA−/−, we report as decreased nuclear circularity, is consistent with many other studies (Lammerding *et al*., 2006; Robijns *et al*., 2016; Chen *et al*., 2018). Thus, while LMNA−/− presents a different phenotype from wild type, VPA, and LMNB1−/−, this different overall loss of shape is consistently reported across other publications in both MEFs and other cell lines. Another study in MEFs reported that loss of all lamins resulted in no nuclear blebs but overall loss of nuclear shape and many nuclear ruptures (Chen *et al*., 2018). Interestingly, shape was restored after adding back lamin A and rupture was suppressed by adding back lamin B1. LMNA−/− cells plus VPA treatment results in increased nuclear blebbing and bleb-based ruptures (**Figure 6**), a phenotype reliant on actin contraction. LMNA−/− nuclei are reported to have disrupted nuclear-actin attachments, specifically due to lamin A/C’s role interacting with SUN1/2 (Broers *et al*., 2004; Chen *et al*., 2018; Vahabikashi *et al*., 2022), which could disrupt actin-contraction-based nuclear bleb formation. Recently, the ability to separate lamin A and lamin C has gained new tools and insights (Wong *et al*., 2021; Vahabikashi *et al*., 2022) which will be vital to future studies needed to better understand the roles of each lamin and chromatin histone modification state. Overall, loss of both lamin A and C provides a different phenotype that may provide the key to how the chromatin and lamins resist actin to maintain nuclear shape and stability.

### Nuclear shape and not ruptures determines increased DNA damage in nuclei

Previous studies have worked to determine the role of both nuclear deformation and ruptures to increased DNA damage (reviewed in (Miroshnikova and Wickström, 2022)). One prominent idea is that DNA damage occurs due to nuclear ruptures because of loss of nuclear repair proteins from the nucleus (Xia *et al*., 2018) and allowing cytoplasmic DNA cutting enzyme TREX1 into the nucleus (Nader *et al*., 2021). However, these studies and many others require the nucleus to transit a confined pore/space resulting in plastic deformation of the nucleus and rupture – an event where drastic shape change, and rupture are interwind. Alternatively, our data clearly show that DNA damage correlates to shape more than rupture (**Figure 4**). Live cell imaging of a nucleus undergoing confined migration shows that the nuclear deformation before rupture induces DNA damage foci formation (Denais *et al*., 2016). Furthermore, deformation without rupture can also increase DNA damage levels (Shah *et al*., 2021). Thus, it is possible that nuclear deformations segregate DNA repair factors away from sites in need of repair (Irianto *et al*., 2017), without necessarily requiring rupture. Our data of nuclear blebbing or decreased nuclear circularity reveal that loss of shape clearly correlates with increased DNA damage. However, increased nuclear ruptures caused by actin contraction activation via Rho activator II (CN03) do not significantly increase DNA damage levels. Thus, our data provides support that nuclear shape deformations drive increased DNA damage, and that increases in nuclear rupture frequency do not.

### Conclusion

Nuclear rupture dynamics studies seeking to understand the role of actin provides insight into an important contributor to disease, loss of nuclear shape and compartmentalization, which then causes nuclear dysfunction. Past studies have focused on the repair of nuclear ruptures and ruptures in confined migration (Denais *et al*., 2016; Raab *et al*., 2016; Halfmann *et al*., 2019; Young *et al*., 2020; Sears and Roux, 2022). However, our data on the dynamics of nuclear blebbing and ruptures in non-migrating and non-artificially confined cells provides insights that help clarify lessons learned from both approaches. One of those main lessons is that there is a clear need to update the model for how actin deforms and ruptures the nucleus. First, actin contraction should now be included as a major antagonist in both unconfined and confined cells. Second, chromatin’s contribution to nuclear shape and rupture must be included, as it is a major mechanical component (Pajerowski *et al*., 2007; Krause *et al*., 2013; Schreiner *et al*., 2015; Shimamoto *et al*., 2017; Stephens *et al*., 2017; Melters *et al*., 2019; Hobson *et al*., 2020; Nava *et al*., 2020; Strickfaden *et al*., 2020). Here we show that chromatin changes are both possibly responsible for some lamin perturbation phenotypes (example LMNB1−/−) and an enhancement of the underlying wild type of behavior. Past schematics and theoretical models of nuclear rupture only account for lamin (lamin A) behavior, which we now know presents differently than wild type, chromatin perturbations, and loss of lamin B1. The ability to incorporate these ideas into both base and theoretical models will aid further investigations aimed at understanding the mechanisms that can disrupt nuclear stability. The relationship between antagonistic external/cytoskeleton forces and the resistive nuclear components to maintain nuclear shape and stability is essential to both basic cell biology and human diseases presenting abnormalities.

## Materials and Methods

### Cell culture and drug treatments

MEF WT, MEF LMNB1−/−, and MEF LMNA−/−, were cultured in DMEM (Corning) containing 10% fetal bovine serum (FBS, HyClone) and 1% penicillin/streptomycin (Corning). The cells were incubated at 37°C and 5% CO2, passaged every 2-3 days and kept for no more than 30 generations.

To treat with drugs, the cells were first plated in DMEM complete and incubated overnight. Cells were then treated with 4 mM VPA (1069-66-5, Sigma), 10 μM Y27632 (129830-38-2, Tocris), 10 nM of Rho Activator II CN03 (Cytoskeleton, Inc), or 2 μM of CytoD (22144-77-0, Tocris). Cells were imaged after 12-24 hours of treatment with VPA and Y27, 3 hours of treatment with CN03, and 1 hour of treatment with CytoD. Cells were not serum starved prior to treatment with CN03.

### Immunofluorescence

Cells were grown in 8 well cover glass chambers (Cellvis) and treated as above. After reaching 80% confluency, cells were fixed with 4% paraformaldehyde (Electron Microscopy Sciences) in PBS (Corning) at room temperature for 15 minutes. The cells were then washed three times with PBS, 5 minutes per wash. After fixation, cells were permeabilized with 0.1% Triton X-100 (US Biological) with PBS for 15 minutes at room temperature. The cells were then washed with 0.06% Tween 20 (US Biological) in PBS for 5 minutes. Cells were washed two more times with PBS, 5 minutes per wash. Cells were blocked in 10% goat serum (Sigma-Aldrich) with PBS for 1 hour at room temperature.

Primary, secondary, and conjugate antibodies were all diluted using the blocking solution (10% goat serum in PBS, Sigma). The primary antibodies used were γMLC2 rabbit Ab 1:100 (3672, Cell Signaling Technologies) and H3K27me^3^ 1:100 (9733, Cell Signaling Technologies). Primary antibodies were added to the dish for 12 hours at 4°C. The cells were then washed three times with PBS for 5 minutes. The secondary antibody used was Alexa Fluor 647 Anti-Rabbit IgG 1:1000 (4414, Cell Signaling Technologies). Secondary antibodies were added to the dish and left to sit at room temperature for 1 hour. Afterwards, the cells were washed with PBS three times. The conjugate antibody used was γH2AX-647 rabbit mAb 1:300 (9720, Cell Signaling Technologies), treated and washed as written above for the primaries.

Next, the cells were stained with a 1 μg/mL dilution of Hoechst 33342 (Life Technologies) in PBS for 5 minutes and then washed with PBS 3 times. The dish was then mounted using ProLong Gold antifade (Life Technologies) and allowed to cure for 12 hours at room temperature.

### Imaging

Images were acquired with Nikon Elements software on a Nikon Instruments Ti2-E microscope with Crest V3 Spinning Disk Confocal, Orca Fusion Gen III camera, Lumencor Aura III light engine, TMC CLeanBench air table, with 40x air objective (N.A 0.75, W.D. 0.66, MRH00401) or Plan Apochromat Lambda 100x Oil Immersion Objective Lens (N.A. 1.45, W.D. 0.13mm, F.O.V. 25mm, MRD71970). Live cell time lapse imaging was possible using Nikon Perfect Focus System and Okolab heat, humidity, and CO2 stage top incubator (H301). Images were captured via camera 16 bit for population images or 12 bit sensitive for time lapse live cell imaging with 40x air objective N.A 0.75 (Nikon MRH00401). Cells were imaged in either 4 well cover glass dishes or 8 well cover glass chambers (Cellvis). For time lapse data, images were taken in 2-minute intervals during 3 hours with 6 fields of view for each condition single plane. Immunofluorescence images were acquired with the 40x air objective at 0.5 μm z-steps over 4.5 μm (9 steps) and maximum intensity compiled post acquisition. Nuclear height measurements were captured using the 100X oil objective using 0.2 μm z-steps over 15 μm.

### γMLC2 analysis

Z-stacks were compiled into a maximum projection and background intensity was measured using a 30 x 30-pixel area containing no cells. A 30 x 30-pixel ROI was drawn around the cell to capture the average intensity of Cy5 fluorescence, then exported from the NIS-Elements software to Excel. Background intensity was subtracted to determine the average levels of yMLC2. Statistical significance was determined using the *t* test.

### DNA damage foci analysis

Z-stacks were compiled into a maximum projection and average background fluorescence was subtracted using a 30 × 30-pixel area containing no cells. Individual nuclei were selected using the NIS-Elements threshold or hand drawn over Hoechst fluorescent images if auto selection was unable to separate between adjacent nuclei. Circularity measurements were taken from auto selected and hand drawn ROIs, then exported from the NIS-Elements software to Excel. We use the bright spots program in Nikon Elements that requires a size and contrast to determine what is a focus. The size of the diffraction limited focus selected was 0.5 μm based on full-width half-maximum experimental measurement of a gaussian from a 175nm fluorescent bead and used across all conditions and biological replicates. Contrast was determined empirically in WT cells and kept constant between treatments in the same replicate. Using the bright spots program Nikon Elements output the number of DNA damage foci per nucleus which was marked by the selected ROI. This data was then exported to Excel for averages and statistical significance determined using the *t* test.

### H3K27me^3^ analysis

As described above, Z-stacks were compiled into a maximum projection and average background fluorescence was subtracted using a 30 × 30-pixel area containing no cells. ROIs were drawn around individual nuclei by hand or using the NIS-Elements threshold. Average intensity of the nucleus was used to determine relative levels of heterochromatin between conditions. Statistical significance was determined using the *t* test.

### Live cell NLS-GFP imaging and analysis

Images were captured via camera 12 bit sensitive with 40x air objective N.A 0.75. Time lapse imaging parameters used were FITC Wide Field light modality at 4% power, 30 ms exposure time at 2-minute intervals during 3 hours with 6 adjacent fields of view for each condition. Images were saved within the NIS-Elements AR Analysis software. Images were observed to record total number of nuclei, number of blebs, and number of ruptures in each field of view. Number of ruptures recorded included each nuclear rupture observed, whether the rupture was bleb-based or non-bleb based, and how frequently each nucleus ruptured throughout the 3-hour duration. Data collected was then compiled and averaged in Excel (Microsoft) to determine percent blebbing, percent rupture, percent bleb-based rupture, and rupture frequency for each condition. Statistical significance was determined by conducting *t* tests between the two conditions.

### Live cell imaging of nuclear height and analysis

Cells were grown in dishes divided into 4 glass wells (Cellvis) and treated as above. After reaching 80% confluency, the cells were treated with 1 μg/mL dilution of Hoechst 33342 (Life Technologies) for 10 minutes before being imaged on a wide-field microscope. For nuclear height measurements, cells were treated with a 1:1000 dilution of SPY555-Actin Probe (CY-SC202, Cytoskeleton) in complete DMEM at 37°C for three hours prior to imaging. Live cell images were taken on a Nikon Eclipse Ti2 wide-field microscope using a 100x oil objective. Image stacks with 0.2 μm steps were also taken using a spinning disc confocal microscope with a 100x oil objective.

Exposure times for Hoechst (DAPI), NLS-GFP (FITC), and SPY555-Actin Probe (TRITC) were between 30 – 100 ms. To determine nuclear height in Z, intensity line scans were taken in Hoechst fluorescence, values were exported to excel and relative positions of the top and bottom of the nucleus were determined using the full-width half-max of the intensity graph in Excel. Two line scan measurements were taken per nucleus and the two values were averaged to determine the height. Statistical significance was determined using the *t* test.

## Acknowledgements

We would like to thank Edward J. Banigan for helpful and insightful discussions and Jeri Ruth Moskowitz for editing support. MP, YB, AG, AL, MLC, KC, AP, ADS are supported by the Pathway to Independence Award (R00GM123195) and Center for 3D Structure and Physics of the Genome 4DN2 grant (1UM1HG011536). The authors declare not competing interests.

---

**Supplemental Figure 1.**
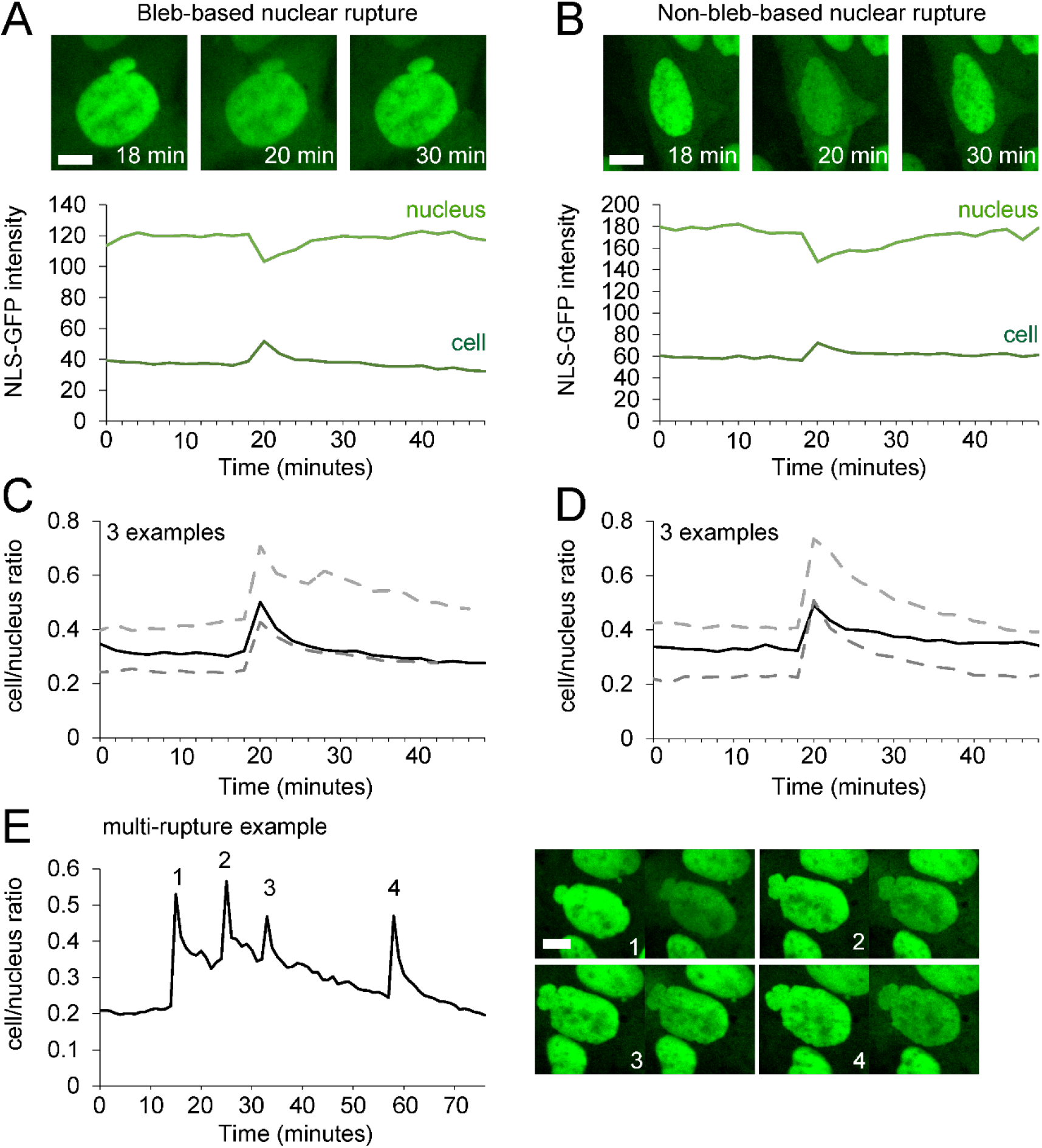
Bleb-based and non-bleb-based nuclear ruptures tracked via nuclear localization signal green fluorescent protein (NLS-GFP). Representative NLS-GFP example images and intensity graph of (A) bleb-based and (B) non-bleb-based nuclear ruptures. Respective graphs show background subtracted nucleus (light green) and cell (dark green) intensities over 48 minutes (24, 2-minute intervals) where rupture occurs at timepoint 20 minutes. While the decrease in nucleus intensity and increase in cell intensity are visible, the ratio of cell/nucleus ratio provides an amplification of nuclear rupture changes for both (C) bleb-based and (D) non-bleb based nuclear ruptures. The black line is the example above (A and B) with two other examples (gray dashed lines). All increases shown are greater than 50% increase in cell/nucleus ratio. From these graphs we quantitatively define nuclear rupture as a > 25% increase in the cell/nucleus NLS-GFP intensity ratio. (E) Multi-rupture bleb-based example cell/nucleus ration graph and images before and after each rupture. Scale bar = 10 μm. See full movies of panel A, B, E with are respectively movies 1, 2, and 3.

**Supplemental Figure 2.**
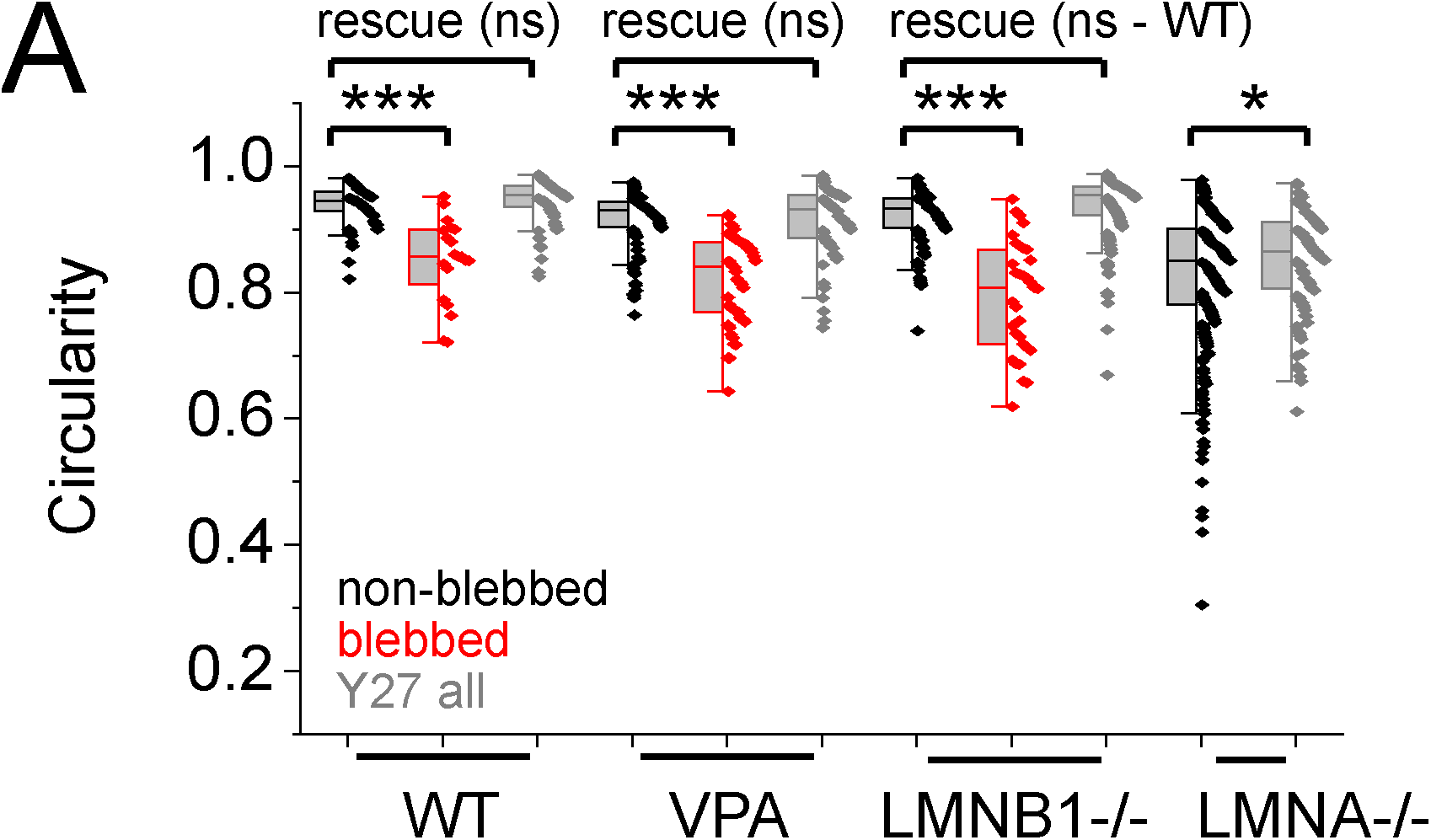
Nuclear circularity provides a measure of abnormal shape. (A) Graph of nuclear circularity for untreated non-blebbed (black) and blebbed nuclei along with all Y27632-treated nuclei (gray) in wild type (WT), chromatin decompaction (VPA), lamin B1 null (LMNB1−/−), and lamin A/C null (LMNA−/−). Individual events are shown as dots relative to box average and 25-75% range. WT, non-blebbed n = 115, blebbed n = 20, Y27632 n = 98; VPA non-blebbed n = 119, blebbed n = 56, Y27632 n = 57; LMNB1−/−, non-blebbed n = 104, blebbed n = 33, Y27632 n = 136; LMNA−/−, non-blebbed n = 287, Y27632 n = 87). (B) Graph of percentage of total nuclei having < 0.9 (black) or < 0.08 (gray). Student’s t-test p values reported as *<0.05, **<0.01, ***<0.001, no asterisk or ns denotes no significance, p>0.05.

**Supplemental Figure 3.**
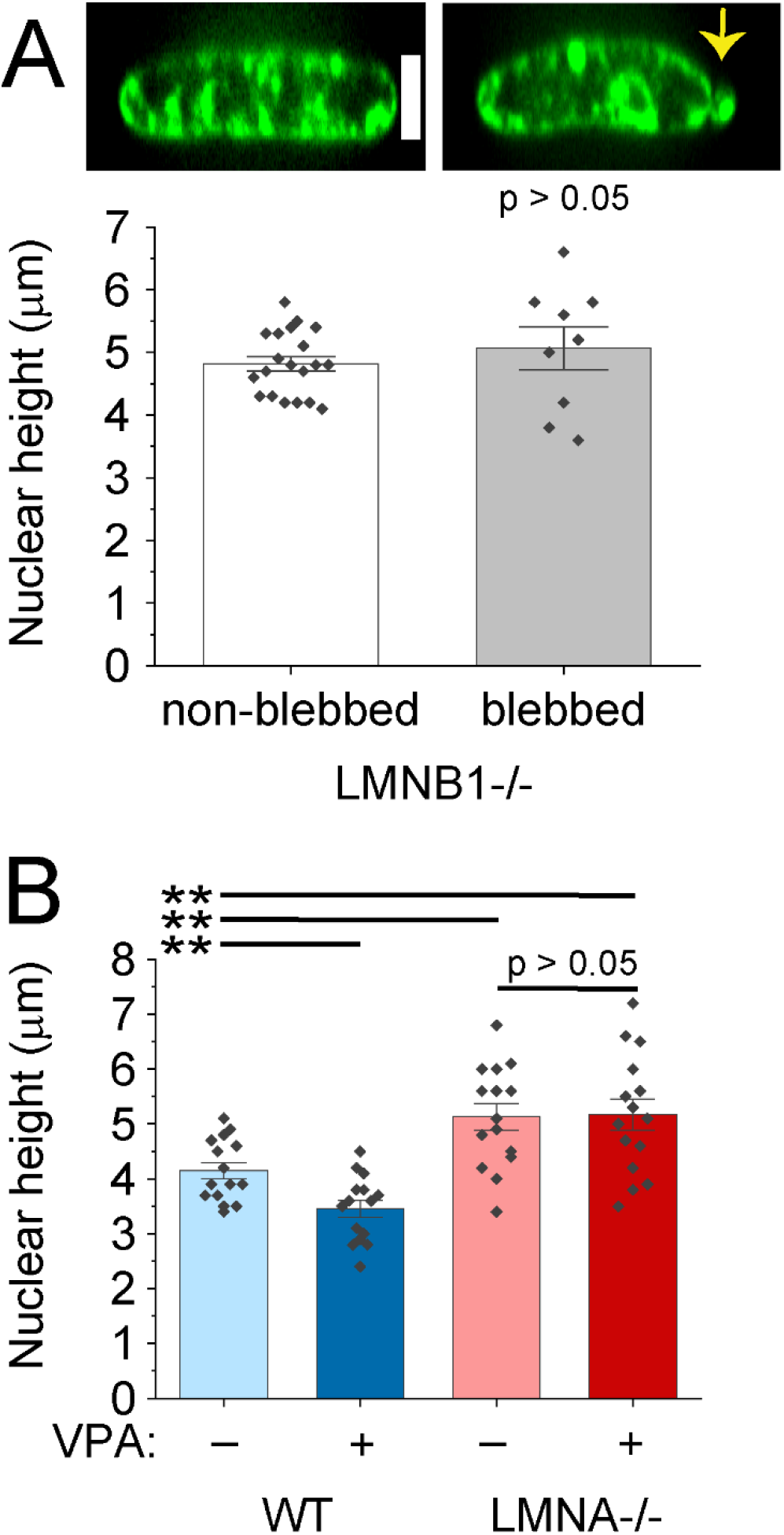
Nuclear height does not change between normal vs. blebbed LMNB1−/− nuclei or upon VPA increased nuclear blebbing in LMNA−/−. (A) Example images and graph of nuclear height measurements of MEF LMNB1−/− non-blebbed (n = 20) and blebbed nuclei (n = 9), where individual measurements are represented as diamonds. (B) Graph of nuclear height for MEF wild type (WT) and lamin A/C null (LMNA−/−) −/+ VPA treatment (n = 15 nuclei shown as diamonds for each). Student’s t-test p values reported as **< 0.01 or no significance p > 0.05. Error bars represent standard error. Scale bar = 5 μm.

**Supplemental Table 1. Raw data.** This excel doc is a compiled document of all raw numbers used in all figures throughout the paper.

